# Male gonad-enriched microRNAs function to control sperm production in *C. elegans*

**DOI:** 10.1101/2023.10.10.561762

**Authors:** Lu Lu, Allison L. Abbott

## Abstract

Germ cell development and gamete production in animals require small RNA pathways. While studies indicate that microRNAs (miRNAs) are necessary for normal sperm production and function, the specific roles for individual miRNAs are largely unknown. Here, we use small RNA sequencing of dissected gonads and functional analysis of new loss of function alleles to identify functions for miRNAs in the control of fecundity and sperm production in *Caenorhabditis elegans* males and hermaphrodites. We describe a set of 29 male gonad-enriched miRNAs and identify a set of 3 individual miRNAs (*mir-58.1, mir-83,* and *mir-235)* and a miRNA cluster (*mir-4807-4810.1)* that are required for optimal sperm production at 20°C and 5 additional miRNAs (*mir-49, mir-57, mir-261,* and *mir-357/358*) that are required for sperm production at 25°C. We observed defects in meiotic progression in *mir-58.1, mir-83, mir-235,* and *mir-4807-4810.1* mutants that may contribute to the reduced number of sperm. Further, analysis of multiple mutants of these miRNAs suggested complex genetic interactions between these miRNAs for sperm production. This study provides insights on the regulatory roles of miRNAs that promote optimal sperm production and fecundity in males and hermaphrodites.

**Article Summary:** MicroRNAs are small non-coding RNAs that are required for the normal production of sperm but the roles of individual microRNAs in the process of spermatogenesis are not well understood. Here, we use the nematode *Caenorhabditis elegans* to identify microRNAs that are enriched in the male gonad to identify specific microRNAs that regulate male fertility. We generated new loss of function mutants for functional analysis to identify a set of microRNAs that are necessary for optimal fertility and fecundity in males.

## Introduction

The production of mature, motile sperm is essential for male fertility in animals. Spermatogonial stem cells undergo mitosis to maintain a pool of progenitor cells capable of entering meiosis and creating haploid spermatids (Handel and Schimenti 2010). Haploid spermatids then differentiate to become motile sperm, which can successfully fertilize an egg. Both mitosis and meiosis are regulated at multiple levels in the male germline to allow for the maintenance and proliferation of the mitotic stem cells, as well as the initiation, progression, and completion of meiosis (Gunes *et al*. 2018). Both transcriptional and translational regulation of gene expression are required to optimally produce sperm (Bettegowda and Wilkinson 2010).

Small RNAs are an important class of post-transcriptional regulators in the process of sperm production (He *et al*. 2009; Papaioannou and Nef 2010; McIver *et al*. 2012; Yadav and Kotaja 2014; Robles *et al*. 2017; Santiago *et al*. 2021). In *C. elegans*, there are four types of endogenous small RNAs, microRNAs (miRNAs), piRNAs, 22G RNAs, and 26G RNAs, that function as guide molecules for Argonaute proteins (Ketting and Cochella 2021). While all of these types of small RNAs regulate gene expression in the germline, the focus of this work is on miRNAs. miRNAs are 21-23 nucleotide, non-coding RNAs that typically bind to the 3’UTR of their mRNA targets with imperfect complementarity. This can result in repression of mRNA translation associated with a destabilization of the mRNA and downregulation of target protein levels (Fabian *et al*. 2010; Jonas and Izaurralde 2015; Chandra *et al*. 2017; Ketting and Cochella 2021).

While there are differences in *C. elegans* spermatogenesis, notably that mature sperm are amoeboid, lacking the flagella common to sperm in most other species, much of the regulation of spermatogenesis is conserved across the animal kingdom (L’Hernault 2009; Ellis and Stanfield 2014). In mammals, miRNA-mediated repression of targets is important for the process of spermatogenesis (Chen and Han 2023). The disruption of miRNA biogenesis is associated with male fertility defects (Hayashi *et al*. 2008; Maatouk *et al*. 2008; Pavelec *et al*. 2009; Korhonen *et al*. 2011; Romero *et al*. 2011; Wu *et al*. 2012; Zimmermann *et al*. 2014; Hilz *et al*. 2016). In humans, expression profiles of testicular and seminal plasma miRNAs are altered in individuals that display sperm defects and infertility, including infertility due to non-obstructive azoospermia (Lian *et al*. 2009; Wang *et al*. 2011; Wu *et al*. 2012, 2013; Abu-Halima *et al*. 2013, 2014; McCubbin *et al*. 2017; Zhang *et al*. 2020; Abu-Halima *et al*. 2021). Additionally, human spermatogenic cells isolated at different stages of spermatogenesis have distinct profiles of miRNAs, suggesting their active involvement in regulating the process of spermatogenesis (Liu *et al*. 2015). While it is clear that miRNAs are necessary for male fertility in mammals, the roles of few individual miRNAs in regulating spermatogenesis have been described (Chen and Han 2023; Hilz *et al*. 2016; Kotaja 2014; Walker 2022).

In *C. elegans*, the function of miRNAs has been shown to be necessary for normal germ cell proliferation and development (Bukhari *et al*. 2012; Dallaire and Simard 2016; Diag *et al*. 2018; Minogue *et al*. 2018, 2020). miRNAs have been identified in mature sperm (Stoeckius *et al*. 2014) and in the dissected gonads of *C. elegans* adults (Minogue *et al*. 2018; Bezler *et al*. 2019). Two miRNA families have been shown to function to regulate sperm production: the *mir-35* family is necessary for male fertility and optimal hermaphrodite sperm production (McJunkin and Ambros 2014) while the *mir-44* family is required for the timing of sperm fate specification in hermaphrodites (Maniates *et al*. 2021).

To identify new functions for miRNAs in the regulation of male fertility and specifically in the process of sperm production and maturation, we performed small RNA sequencing to identify miRNAs in isolated gonad arms from males and hermaphrodites. Concurrent with this small RNA sequencing work, a similar study by Bezler *et al* (2019) was published. That study provided a comprehensive analysis of all classes of small RNAs in hermaphrodites and males and identified sex differences in response to environmental RNAi, whereas the focus of this study is the functional analysis of male gonad-enriched miRNAs. In order to determine whether male gonad-enriched miRNAs function to regulate male fertility, new loss of function mutants were generated for a set of 29 male gonad-enriched miRNAs. Mutants were analyzed to identify defects in male fertility. New functions were identified for male gonad-enriched miRNAs, including in the regulation of male mating, sperm production, and meiotic progression. Two miRNAs (*mir-58.1,* and *mir-235)* function to regulate sperm production in both males and hermaphrodites, while one miRNA (*mir-83)* and one miRNA cluster (*mir-4807-4810.1)* functions to control sperm production specifically in males. Loss of these three miRNAs or one miRNA cluster is associated with defects in meiotic progression in the male germline. Lastly, genetic analysis indicates complex interactions between male gonad-enriched miRNAs and likely opposing activities of these miRNAs, though the specific mechanism of action is not yet known.

## Results

### Identification of miRNAs in male and hermaphrodite gonads

To investigate miRNA regulation of sperm production and function, we sought to identify miRNAs that are present in male and hermaphrodite gonad arms. In *C. elegans*, germ cells are located in gonad arms, in which they undergo mitotic proliferation at the distal ends of the arm with meiotic maturation and differentiation in the more proximal regions to produce functional sperm and oocytes. Hermaphrodite gonad arms produce sperm during the last larval stage (L4) and then switch to oogenesis, while the male gonad arm produces sperm continuously starting in the L4 stage. Therefore, adult male and adult hermaphrodite gonads are spermatogenic and oogenic, respectively (Figure 1A). By comparing miRNAs expressed in spermatogenic and oogenic gonad arms, miRNAs that regulate sperm production and function could be revealed. We dissected gonad tissue from adult males and adult hermaphrodites for small RNA sequencing (Figure 1B) and identified 181 out of the total 253 miRNAs from miRBase Release 22 (Figure S1, Table S1). There was about 82% overlap in the miRNAs found in both male and hermaphrodite gonads, with 148 shared miRNAs (Figure 1C). A subset of 14 miRNAs were found only in males while 19 were only found in hermaphrodites (Figure 1C). Differential expression analysis between male and hermaphrodite gonad miRNA profiles found 29 miRNAs that had higher expression levels in male gonads and 32 that had higher expression levels in hermaphrodite gonads with a fold change >2 and p-value <0.01 (Figure 1D). Another study also identified miRNAs in isolated gonads from males and hermaphrodites (Bezler *et al*. 2019). We compared the miRNA profiles from both studies and found that the miRNAs identified largely overlapped in both hermaphrodite (Figure 1E, 133 miRNAs) and male gonads (Figure 1F, 144 miRNAs).

**Figure 1.**
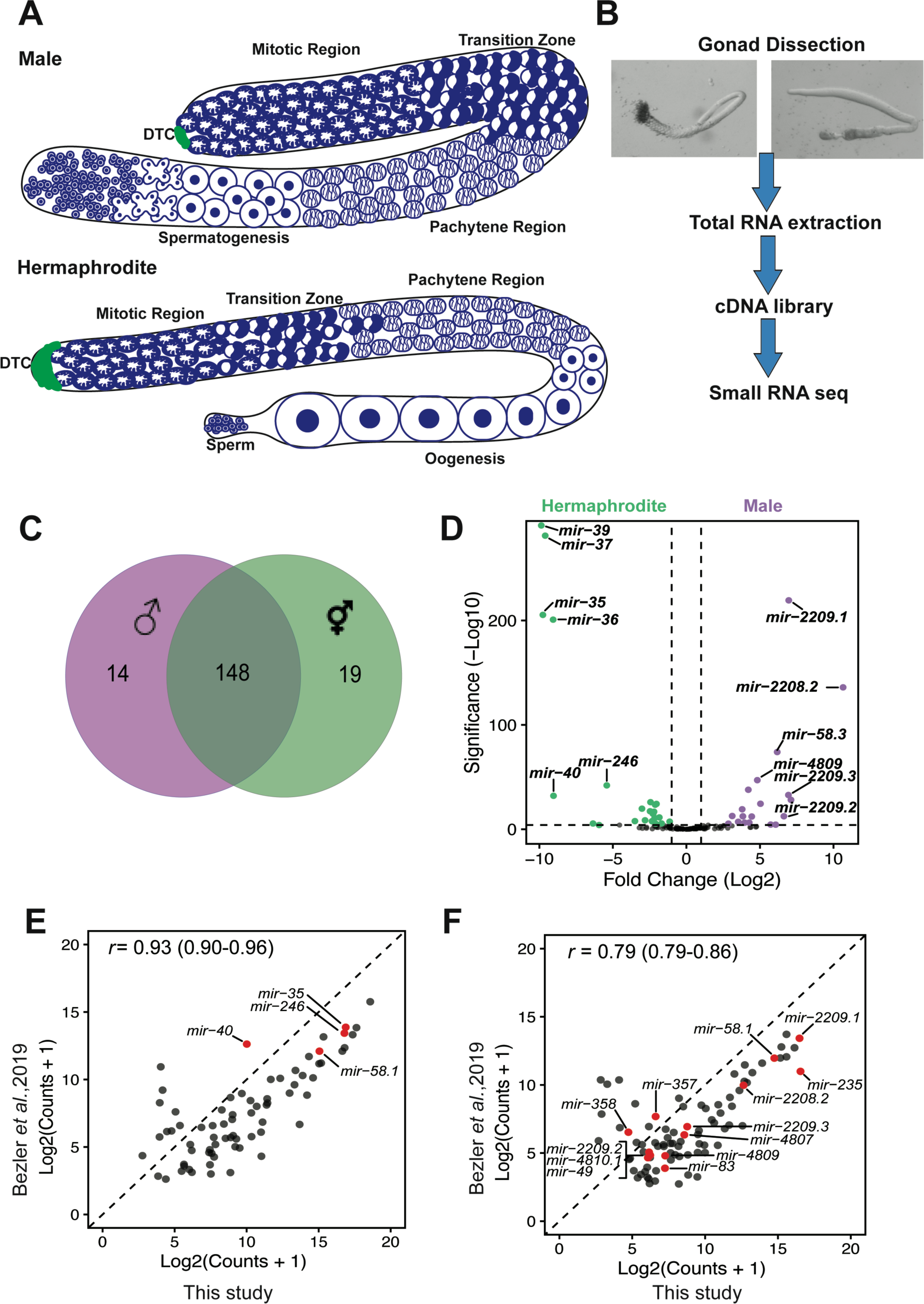
Differential miRNA profiles in isolated male and hermaphrodite gonads. (A) Cartoon of adult male (spermatogenic) and hermaphrodite (oogenic) gonad showing the appearance of chromatin in nuclei in the mitotic and meiotic regions. Only half of the gonad was shown for hermaphrodite. (B) Procedure for gonad isolation for small RNA sequencing analysis. (C) Gonad expression of miRNAs in male (purple) and hermaphrodites (green). Total number of miRNAs in each sex and the overlap between the sexes are shown. (D) Differential expression analysis of miRNA profiles between males and hermaphrodites. With log2(fold change)>1 and Padj<0.01, male and hermaphrodite gonad-enriched miRNAs are highlighted in purple, and green, respectively. (E) The gonad-enriched miRNA profile shared between this study(x-axis) and Bezler *et al* (2019).(y-axis), for hermaphrodite gonads (E) and male gonads (F). Only miRNAs detected with a mean normalized count>= 5 were included. The Pearson correlation coefficient with 95% confidence intervals is shown in the top left corner.

### Creation of miRNA loss of function mutants for functional analysis

To test whether the 29 miRNAs that were found to be enriched in male gonads function to regulate spermatogenesis, we sought to perform functional analysis on miRNA loss of function mutants. However, only 8 of the 29 miRNAs had existing deletion mutants available: *mir-49, mir-57, mir-75, mir-83, mir-235, mir-261, mir-357,* and *mir-358* (Table 1). We also included *mir-58.1* in our analysis. Of the 21 remaining male gonad-enriched miRNAs, 13 are located in two genomic clusters, the *mir-2209.2-mir-2209.3* cluster on chromosome IV (*mir-2209.2, mir-2208.1, mir-2208.2, mir-2209.1, and mir-2209.3)* (Figure 2A) and the *mir-4807-mir-4923.1* cluster on the X chromosome (*mir-4807, mir-4808, mir-4809, mir-2220, mir-4810.2, mir-4810.1, mir-1018, mir-4925, mir-4923.2,* and *mir-4923.1* (Figure 2B). The *mir-4807-4923.1* cluster comprises two smaller clusters, *mir-4807-4810.1* and *mir-1018-4923.1* (Figure 2B). The sequence of the *mir-4807-4923.1* cluster is located within the genomic sequence for an uncharacterized non-coding RNA, *Y59E1B.1* (Figure 2B). Therefore, the set of deletion mutations for miRNAs in this cluster also disrupts the sequence of *Y59E1B.1*. In order to perform functional analysis on the full set of 29 male gonad-enriched miRNAs, we generated new loss of function mutants missing either single or multiple clustered miRNAs using CRISPR-Cas9 genome editing. A total of 16 new miRNA loss of function alleles were created (Table 1), backcrossed, and analyzed for defects in hermaphrodite and male fertility. To facilitate phenotypic analysis, strains were constructed with *him-8(e1489)* for generation of males and a *his-72::gfp* transgene (*stIs10027*) (Huang *et al*. 2012) for quantification of GFP positive mature spermatids. Analysis was performed in this genetic background unless otherwise noted.

**Figure 2.**
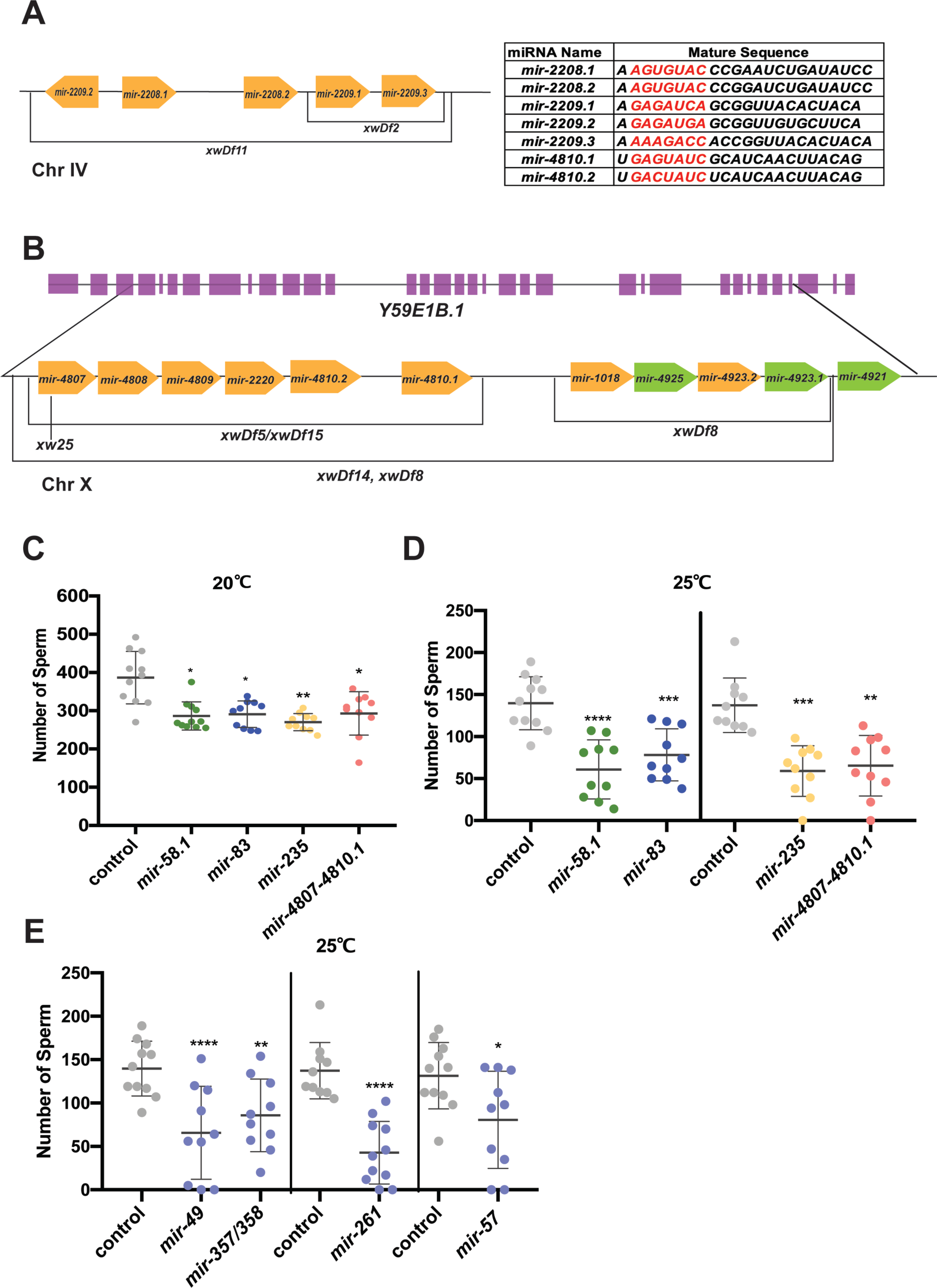
Subset of miRNA mutant males produced fewer spermatids at 20°C and 25°C. (A) (B) Cartoon showing two miRNA clusters containing 13 out of 29 male gonad-enriched miRNAs (yellow) and the loss of function alleles generated. The table on the top right corner shows the mature sequence of some gonad enriched miRNAs that share a seed sequence. The relative genomic location was not scaled. (A) *mir-2209.1-mir-2209.3* cluster on chromosome IV. (B) The genomic location of *mir-4807-4023.1* on chromosome X relative to Y59E1B.1. The miRNAs highlighted in green are not significantly expressed between male and hermaphrodite gonads. (C-D) The number of sperm for individual worms was quantified using *his-72::gfp* expression to detect the condensed chromatin of haploid spermatids. All strains assayed have *him-8(e1489); stls10027 (Phis-72::HIS-72::GFP) (him-8; his-72)*. The scatter plots show the number of spermatids produced in control (*him-8; his-72,* gray) and mutant males with each dot representing an individual worm and the lines representing mean ± SD. The statistical test between control and mutant strains were performed using Dunnett’s or Dunnett’s T3, and the significant difference is represented as *, p<0.05; **, p<0.01; ***, p<0.001; and ****, p<0.0001 above each data set. The results were listed in Table S2. Males were counted at L4 molt + 5 hours and L4 molt at 20°C and 25°C, respectively. (C) Number of male sperm at 20°C. (D) Number of male sperm at 25°C. (E) Number of male sperm at 25°C.

**Table 1.** Identification of Male Gonad-enriched miRNAs.

| miRNA | Base Mean<sup>a</sup> | log<sub>2</sub>(FC)<sup>b</sup> | P-adj<sup>c</sup> | Loss of function alleles<sup>d</sup> | Expression pattern described in Bezler *et al.*, (2019)<sup>e</sup> |
| --- | --- | --- | --- | --- | --- |
| *mir-2208.2* | 3193.62 | 10.65 | 1.10e-136 | *xwDf11* | male somatic gonad bias |
| *mir-2208.1* | 109.77 | 7.11 | 7.49e-29 | *xwDf11* | male somatic gonad bias |
| *mir-2209.1* | 46069.77 | 6.96 | 4.72e-220 | *xwDf2*,<br>*xwDf11* | male somatic gonad bias |
| *mir-2209.3* | 219.84 | 6.94 | 1.61e-33 | *xwDf2*,<br>*xwDf11* | male somatic gonad bias |
| *mir-2209.2* | 35.45 | 6.63 | 3.61e-13 | *xwDf11* | male somatic gonad bias |
| *mir-58.3* | 288.70 | 6.17 | 1.06e-74 | *xw39* | not annotated |
| *mir-58.1* | 30679.86 | -0.30 | 0.41 | *n4640* | in males, no sex bias |
| *mir-261* | 5.20 | 6.07 | 4.21e-5 | *n4594* | reads below threshold<sup>f</sup> |
| *mir-789.2* | 59.01 | 5.73 | 3.57e-5 | *xw61* | male somatic gonad bias |
| *mir-4809* | 78.71 | 5.04 | 3.89e-25 | *xwDf5*,<br>*xwDf14*,<br>*xwDf15* | male somatic gonad bias |
| *mir-1018* | 1727.51 | 4.82 | 9.36e-48 | *xwDf18* | male somatic gonad bias |
| *mir-2221* | 3.69 | 4.77 | 2.55e-3 | *xw53* | reads below threshold |
| *mir-4808* | 104.63 | 4.44 | 6.64e-13 | *xwDf5*,<br>*xwDf14*,<br>*xwDf15* | male somatic gonad bias |
| *mir-789.1* | 44.00 | 4.27 | 4.97e-7 | *xw34* | male somatic gonad bias |
| *mir-4807* | 200.20 | 4.21 | 1.48e-38 | *xw25, xwDf5, xwDf14, xwDf15* | male somatic gonad bias |
| *mir-2220* | 104.49 | 3.97 | 4.97e-7 | *xwDf5, xwDf14, xwDf15* | male somatic gonad bias |
| *mir-4810.1* | 39.76 | 3.84 | 4.00e-13 | *xwDf5, xwDf14, xwDf15* | male somatic gonad bias |
| *mir-1822* | 50.29 | 3.79 | 7.50e-20 | *xw45* | in males, no sex bias |
| *mir-4810.2* | 96.69 | 3.52 | 7.75e-8 | *xwDf5, xwDf14, xwDf15* | reads below threshold |
| *mir-4923.2* | 31.60 | 3.11 | 2.05e-13 | *xwDf18* | reads below threshold |
| *mir-2210* | 21.19 | 3.10 | 6.33e-4 | *xw36, xw55* | male bias |
| *mir-1819* | 36.26 | 2.85 | 4.59e-6 | *xw42* | male germline bias |
| *mir-49* | 38.64 | 2.56 | 5.63e-4 | *zen99* | male bias |
| *mir-357* | 56.16 | 2.53 | 2.95e-4 | *nDf60* | male bias |
| *mir-235* | 57278.28 | 2.33 | 2.33e-3 | *n4504* | in males, no sex bias |
| *mir-358* | 15.30 | 2.29 | 2.64e-3 | *nDf60* | male bias |
| *mir-796* | 13.66 | 2.02 | 8.93e-4 | *xw58* | reads below threshold |
| *mir-83* | 102.67 | 1.48 | 1.14e-4 | *n4638* | in males, no sex bias |
| *mir-57* | 25945.94 | 1.27 | 5.84e-3 | *gk175* | in males, no sex bias |
| *mir-75* | 71.11 | 1.15 | 3.48e-4 | *n4472* | in males, no sex bias |

<sup>a</sup> Base mean=the average of the normalized counts in DESeq2.
<sup>b</sup> $\log_2(\text{FC}) = \log_2 \text{Fold Change}$ between male and hermaphrodite gonads in DESeq2. <sup>c</sup>
P-adj, Benjamini-Hochberg adjusted p-value in DESeq2.
<sup>d</sup> all *xw* alleles were generated in this study.
<sup>e</sup> comparison of expression patterns between hermaphrodites, males, and feminized *fog-3* males allowed for the determination of expression bias in males or hermaphrodites, and in somatic gonad or germline Bezler *et al.* (2019)
<sup>f</sup> sense reads in Bezler *et al.* (2019) below minimum threshold of 5 mean sense reads across replicates.

### Three miRNAs necessary for male mating efficiency

We first used a mating assay as a screening tool to identify miRNA mutant males that had defects in mating behavior, sperm production, or sperm function. Single mutant males were analyzed for the ability to mate and produce cross progeny. No gross defects in male morphology or motility were observed. Reduced mating efficiency was found associated with the loss of three miRNAs: *mir-83, mir-789.1,* and *mir-2221* (Figure S2A). We further analyzed these mutant males using a sperm transfer assay in which the interactions between mutant males and control hermaphrodites were observed and the transfer of male sperm was assessed. While the number of interactions between *mir-789.2* and *mir-2221* mutant males and hermaphrodites was comparable to control males, *mir-83* mutant males displayed few interactions with hermaphrodites (Figure S2B). The few interactions that were observed appeared briefer than control male interactions and failed to result in sperm transfer (Figure S2C). Together, this suggests that *mir-83* mutant males may fail to sense or respond to hermaphrodites.

We next wanted to analyze whether *mir-789.2* and *mir-83* functioned redundantly with their respective family members, *mir-789.1* and *mir-49* (Figure S2D), which were also found enriched in male gonads, in the control of male mating. miRNAs that share a seed sequence are grouped into miRNA families and often function redundantly (Miska *et al*. 2007; Alvarez-Saavedra and Horvitz 2010). Surprisingly, males of both double mutants displayed normal mating success (Figure S2E). This suggests possible opposing roles for these miRNAs in the regulation of mating behavior.

Apart from *mir-83* mutant males, which fail to mate with hermaphrodites, the remaining 25 miRNA mutants analyzed showed that, upon successful mating by mutant males, there were a large number of cross-progeny generated with a low percentage of self-progeny comparable to control worms (Table S2). For these strains, male sperm was preferentially used to fertilize the oocytes and successfully produce cross-progeny, comparable to control males. Together, this indicates that male sperm function is not compromised in this set of miRNA mutants.

### miRNAs regulate sperm production

With overall normal male fertility observed in miRNA mutants, we next asked whether there were any defects in the rate of sperm production in this set of miRNA loss of function mutants. While hermaphrodites generate a finite number of sperm during larval development, males produce sperm continuously starting in the L4 larval stage. To analyze sperm production, males were synchronized at the L4 molt stage and sperm were counted at specific times following the L4 molt. Differences in sperm number could result from changes in the timing of sperm onset, the rate of germline mitosis, or the rate of meiotic progression to form the mature haploid spermatids that were counted. Four miRNA mutants, *mir-58.1*, *mir-83, mir-235,* and *mir-4807-4810.1,* were observed to have a lower average number of sperm in young adult males analyzed 5 hours after the L4 molt (Figure 2C and S3A). Although the sperm count in other miRNA mutant males was not affected at 20°C (Table S2), it is possible that these miRNAs function to maintain normal spermatogenesis at an elevated temperature of 25°C, which is the high end of the normal cultivation temperature range of *C. elegans* but provides a moderate temperature stress. To test this, we analyzed the number of sperm in F2 mutant males at the L4 molt stage grown at 25°C after upshifting the P0 worms from 20°C to 25°C. *mir-58.1*, *mir-83, mir-235,* and *mir-4807-4810.1* mutant males showed a further reduction of sperm at 25°C (Figure 2D). An additional 4 miRNA mutants had a lower sperm count at 25°C (Figure 2E) despite having a normal sperm count at 20°C (Figure S3B).

Loss of the *mir-4807-4810.1* cluster was the only male gonad-enriched miRNA cluster mutation (Figure 2B) found to affect sperm production. Loss of the two smaller clusters that comprise the larger *mir-4807.1-mir-4923.1* cluster, *mir-4807-4810.1* and *mir-1018-4923*.1, didn’t result in a reduction in sperm number (Table S2). This suggests that the defects observed in *mir-4807-4810.1* mutants were suppressed by the loss of *mir-1018-4923.1*. The mechanism underlying this suppression is unknown. Surprisingly, although the *mir-2209.2-2209.3* miRNA cluster contains 5 miRNAs with the highest fold change in our differential expression analysis, its loss did not result in a reduced number of sperm (Table S2, Figure S3C). Lastly, loss of all 13 miRNAs found in the two male gonad-enriched clusters in *mir-2209.2-2209.3; mir-4807-4810.1 mir-1018-4923.1* mutants was not associated with a reduction in sperm number (Figure S3C).

We tested for genetic interactions with selected family members of the four miRNAs for which mutants had reduced sperm. First, interactions between *mir-58.1* and *mir-58.3* were analyzed. Unlike its family member *mir-58.3, mir-58.1* doesn’t have higher expression level in male gonads (Table 1). We found no enhancement of the reduced sperm number phenotype in the *mir-58.1 mir-58.3* double mutant males (Figure S3D), but rather saw suppression of the mutant phenotype at 25°C (Figure S3E). Next, loss of *mir-49* suppressed the reduced sperm number phenotype in *mir-83* mutant males at 20°C but enhanced the phenotype at 25°C (Figure S3D, E), suggesting that *mir-49* functions differently under normal and stressed conditions. Together, these results were not consistent with the simple model that miRNA family members function redundantly to regulate shared targets, but rather suggest more complex genetic interactions that affect the male gonad.

### miRNAs regulate fecundity and sperm production in hermaphrodites

Since hermaphrodites produce both sperm and oocytes, defects in sperm production or function can result in reduced hermaphrodite fecundity. To test this, brood size analysis was performed in hermaphrodites for our set of miRNA mutants (Table S3). Seven miRNA mutant strains had a decreased number of progeny compared to control hermaphrodites (Figure 3A). Next, the number of sperm produced by mutant hermaphrodites was determined (Table S4). Three of the seven miRNA mutant strains with reduced brood sizes also had fewer sperm (Figure 3B). Thus, for these three strains, the reduced number of sperm is likely responsible for the observed lower brood size. Interestingly, the brood size associated with loss of *mir-2221* without *him-8; his-72* in the genetic background was not affected compared to control N2 hermaphrodites (Figure 3C), suggesting that the *him-8; his-72::gfp* is a weakly sensitized genetic background.

**Figure 3.**
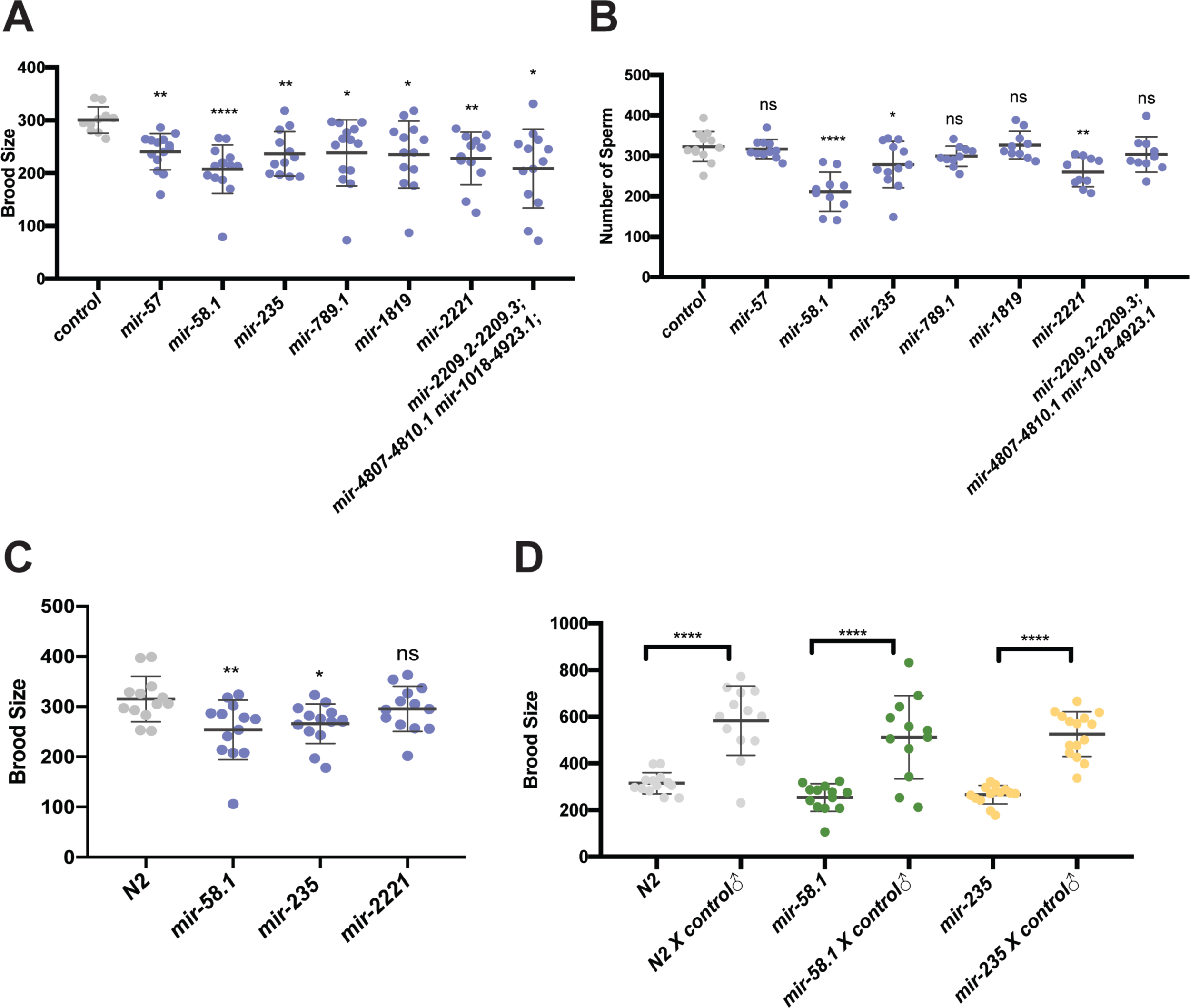
Subset of miRNA mutant hermaphrodites show reduced fecundity and fewer spermatids. Brood size analysis (A,C,D) and sperm quantification (B) was performed on miRNA mutant strains and results from mutant strains are shown compared to control (*him-8; his-72*, gray). Scatter plots show brood size (A,C,D) or sperm number (B) in control and mutant hermaphrodites with each dot representing data from individual worms and the lines representing mean ± SD. The statistical test between control and a specific mutant strain was performed using Dunnett’s or Dunnett’s T3. And the significant difference is represented as *, p<0.05; **, p<0.01; ***, p<0.001; and ****, p<0.0001. The results were listed in Table S3 and S4. (A) Brood size analysis and (B) sperm quantification in the 7 miRNA mutant strains with significantly lower brood sizes compared to control (*him-8; his-72*, gray) are shown. (C) Brood size analysis of 3 miRNA mutant strains with lower sperm number compared to N2 in the absence of *him-8; his-72* in the genetic background. (D) Brood size analysis of unmated and mated miRNA mutant strains along with N2 controls. Welch’s T-test was used to compare brood size between unmated and mated hermaphrodites. Hermaphrodites were mated with *him-8; his-72* males and the number of progeny were counted for individual worms.

To test whether the reduced number of hermaphrodite sperm can account for the reduced brood size, we tested whether mating with control males could restore normal fecundity. The brood sizes of *mir-58.1* and *mir-235* mutant hermaphrodites were increased when mutants were mated with control males (Figure 3D), indicating defects in sperm, not oocyte, production in mutant hermaphrodites. Together, these results indicate that the regulatory roles of *mir-58.1* and *mir-235* in sperm production is shared by males and hermaphrodites, while *mir-83* and *mir-4807-4810.1* is important specifically in sperm production in males.

### Genetic interactions between the set of four male gonad-enriched miRNAs involved in sperm production

To test for genetic interactions between *mir-58.1*, *mir-83, mir-235*, and *mir-4807-4810.1*, we analyzed the number of sperm produced in multiply mutant males (Table S5). Although these four miRNA mutants all showed a lower number of sperm in young adult males, when combined we observed that the defects were not strictly additive (Figure 4A). First, some combinations of miRNA mutants showed no further reduction in sperm number compared to the single mutants. For example, the mutant males with loss of *mir-235* and the *mir-4807-4810.1* cluster had sperm counts comparable to *mir-4807-4810.1* mutants (Figure 4B). Second, some combinations of mutants showed suppression of the reduced number of sperm: *mir-235*; *mir-83* double mutant males displayed sperm counts higher than both single mutants and not statistically different from controls (Figure 4C). Third, we observed enhanced defects: the *mir-83; mir-4807-4810.1*, *mir-58.1; mir-235,* and *mir-58.1; mir-83* double mutant males displayed an enhanced phenotype compared to the respective single mutants (Figure 4D, Figure S4A,B). The male sperm count was further reduced in *mir-235*; *mir-58.1 mir-83* triple mutant (Figure S4C), suggesting that *mir-58.1* acts with or in parallel to *mir-235* and *mir-83*. Together, the evidence from analysis of multiple miRNA mutants suggests a complex genetic network for *mir-235*, *mir-4807-4810.1*, *mir-58.1*, and *mir-83* to allow optimal sperm production.

**Figure 4.**
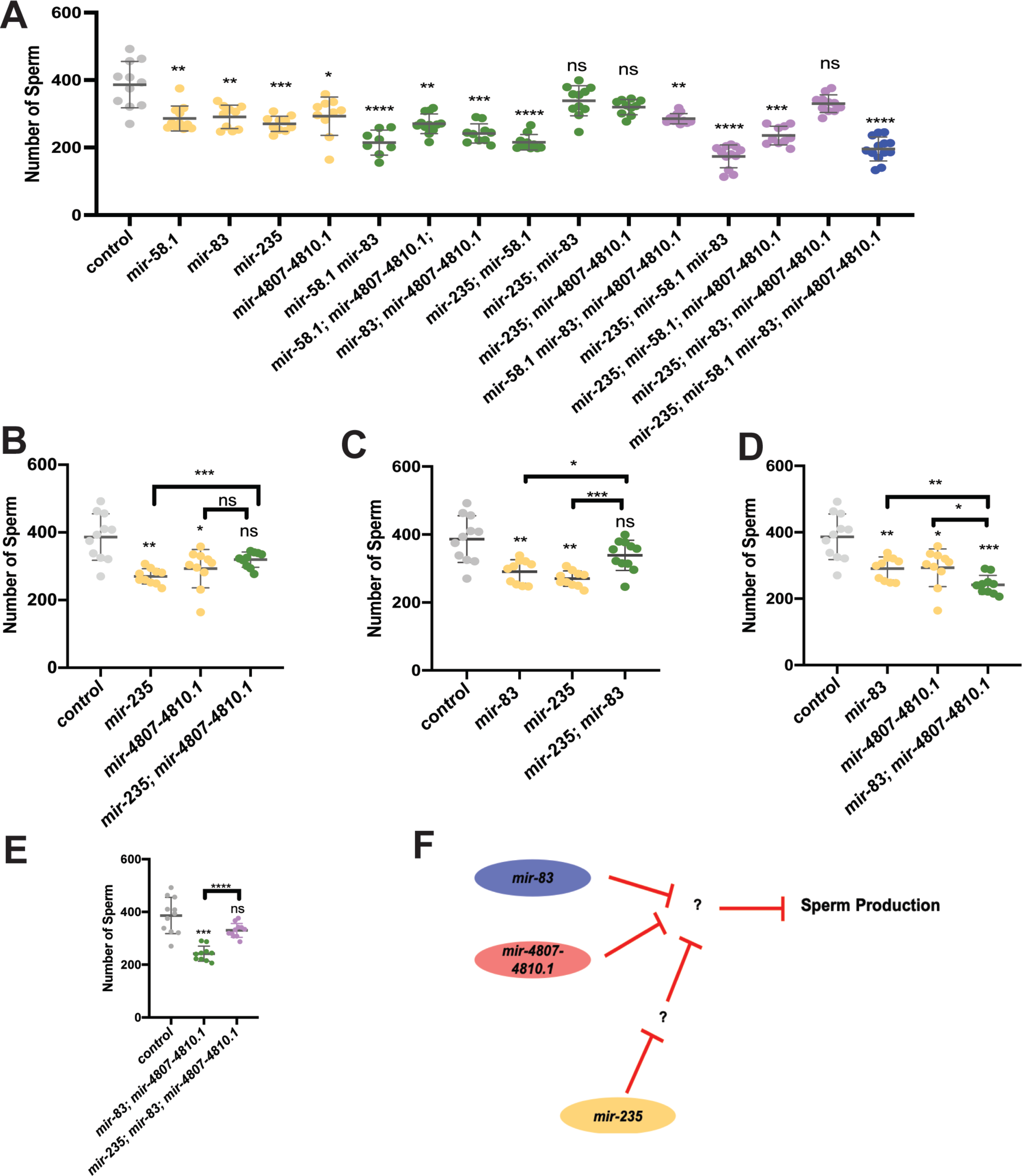
Complex regulatory network of miRNAs controls sperm production. Genetic interactions were analyzed between *mir-58.1, mir-83, mir-235* and *mir-4807-4810.1.* Sperm quantification was performed for individual control and miRNA multiply mutant strains using *his-72::gfp* to detect haploid spermatids. Each dot in the scatter plot represents the number of sperm in individual worm and lines represent mean ± SD for each strain. All strains assayed with *him-8; his-72* in the genetic background. The statistical analysis results are represented as ns, p>0.05; *, p<0.05; **, p<0.01; ***, p<0.001; and ****, p<0.0001. The comparison between control (*him-8; his-72*) and mutant is indicated above the data sets with Dunnett’s T3, and other comparison pairs are indicated by the line above them by Welch’s T-test. The results are listed in Table S5. (A) The number of haploid spermatids in controls (gray), single (yellow), double (green), triple (purple), and quadruple (blue) mutant males. (B-D) Selected datasets from data shown in (A) to highlight interactions in single and double mutants. (E) Genetic interactions analyzed between *mir-235* and *mir-83; mir-4807-4810.1.* (F) Diagram of the genetic interactions of miRNAs that regulate sperm production.

Interestingly, loss of *mir-235* partially or fully suppressed the phenotype in either *mir-83* or *mir-4807-4810.1* (Figure 4B and 4C). And the phenotype in *mir-235; mir-83; mir-4807-4810.1* triple mutant males was suppressed compared to the double (Figure 4E). Together, the results indicate that *mir-235* may act in a mechanism that is antagonistic to *mir-4807-4810.1* and *mir-83* (Figure 4F).

Lastly, we asked whether the genetic interactions observed in sperm count analysis were also observed in brood size analysis in hermaphrodites. Brood size analysis was conducted for multiple miRNA mutants (Figure S5A). The brood size in *mir-235; mir-83* is comparable to control, suggesting that the antagonistic roles between them is shared by both male and hermaphrodite (Figure 4C, Figure S5B). The effect of losing *mir-235* and *mir-58.1* is comparable to losing *mir-58.1* alone (Figure S5C), unlike the additive effect observed in the reduced sperm number in males (Figure S4A). Interestingly, although we didn’t observe an effect of losing *mir-4807-4810.1* on hermaphrodite sperm production and hence no effect on brood size, it was found that the brood size in *mir-58.1; mir-4807-4810.1* double mutant was increased compared to *mir-58.1* (Figure S5D).

### Male gonad-enriched miRNAs necessary for meiotic progression

To determine whether the reduced number of sperm in *mir-58.1*, *mir-83, mir-235*, and *mir-4807-4810.1* mutant males is associated with defects in mitotic or meiotic progression in the male germline, analysis of nuclear morphology was performed using DAPI staining of mutant male gonads (Albert Hubbard and Schedl 2019). In male gonad arms, the germ cells in the meiotic transition zone have distinct, polarized chromatin morphology, which can be easily identified and used to distinguish mitotic cells from early meiotic cells (Figure 1A). In N2 males, there is an average of 27 and 18 rows of nuclei in the mitotic and meiotic transition zone areas, respectively (Morgan *et al*. 2010). Quantification of rows of nuclei in the isolated gonad arms of mutant males indicated that the mitotic region was similar to controls (Figure S6A). However, we observed that the transition zone length was shorter in the gonad arms of *mir-58.1*, *mir-83, mir-235*, and *mir-4807-4810.1* mutants (Figure 5A). No additional morphology defects were observed in the gonad arms of miRNA mutants. While the mitotic zone length remained comparable to controls in the multiply mutant males (Figure S6B), the transition zone length was further shortened in the multiply mutant males that showed enhanced sperm defects (Figure 5B). This correlation suggests that meiotic progression defects may be associated with the lower sperm count in miRNA mutants. The reduced sperm number phenotype in *mir-235; mir-83; mir-4807-4810.1* mutant males was suppressed compared to the double (Figure 4G). However, no such suppression effect was identified for the transition zone length (Figure 5B), suggesting that this suppression occurs downstream or independent of the meiotic progression phenotype.

**Figure 5.**
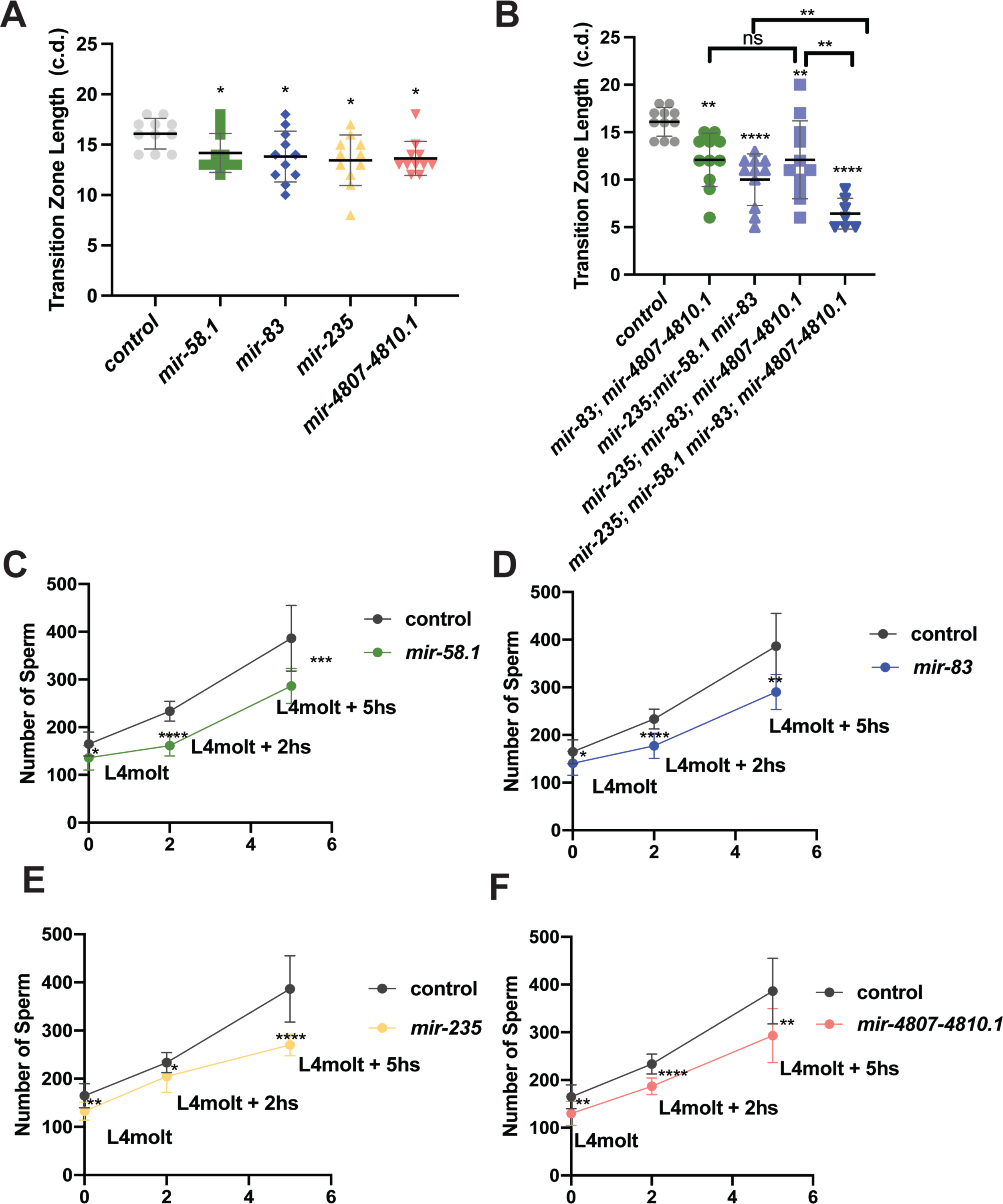
Shorter transition zones observed in miRNA mutants with reduced sperm production. (A-B) Individual DAPI stained gonads were analyzed for the length of the transition zone, determined by the start of nuclei with polarized chromatin in a crescent morphology and the start of pachytene nuclei in miRNA single mutants (A) or multiply mutants (B). Each dot represents the transition zone length of individual gonad in cell diameter (c.d.). Error bar shows the mean ± SD. 7-11 gonads analyzed for all strains. The statistical analysis between control (*him-8; his-72*) and mutant was conducted by Dunnett’s test. The comparison between multiple mutants is indicated by the line above them with Welch’s T-test. (C-F) Sperm quantification in *mir-58.1, mir-83, mir-235* and *mir-4807-4810.1* mutants at three time points. The number of haploid spermatids in control males (gray) and miRNA mutant males at L4 molt, L4 molt+2 hours, and L4 molt+ 5 hours. On each plot, the mean number of sperm was showed as dot with error bars showing mean ± SD. The statistical analysis between control and mutant was conducted by T-test and represented with ns, p>0.05; *, p<0.05; **, p<0.01; ***, p<0.001; and ****, p<0.0001.

The reduction in sperm number in young adult mutant males could be due to a slower rate of sperm production or to a delay in the onset of haploid spermatid production at L4 stage. To examine this, we first quantified male sperm at earlier time points after the L4 molt. The results showed that the mutants had lower male sperm count than control at all time points examined (Figure 5C-5F), which may indicate a slower rate of sperm production. At the L4 molt, the miRNA mutants already displayed lower male sperm count. Therefore, we determined whether the timing of when haploid spermatids start to be produced is affected in these mutants. Early L4 stage males were analyzed for the appearance of haploid spermatids in mutant gonad arms. In *mir-4807-4810.1* mutants, none of the early L4 stage males before the start of tail retraction had sperm production compared to 10% in control (Figure S6C). For early L4 stage males with ongoing tail retraction, 76% of *mir-4807-4810.1* mutant worms showed sperm production compared to 100% of control worms (Figure S6D). This suggests that a delay of spermatid onset may also contribute to the reduced sperm phenotype in *mir-4807-4810.1* mutant males. Taken together, defects in meiotic progression and a slower rate of sperm production may contribute to the lower sperm number in *mir-58.1*, *mir-83, mir-235*, and *mir-4807-4810.1* mutant males.

To test whether meiotic progression is affected in the mutant males, we performed an EdU pulse-chase experiment. To measure meiotic progression, we exposed control and mutant males to an EdU pulse for 2.5 hours followed by a 10-hour chase. In this way, the progression of a set of cells through meiosis can be monitored (Jaramillo-Lambert *et al*. 2007; Kocsisova *et al*. 2018; Almanzar *et al*. 2021; Cahoon and Libuda 2021). We investigated the rate of meiotic progression in the *mir-235; mir-58.1 mir-83 him-8; mir-4807-4810.1* multiply mutant males, which displayed the strongest reduction in sperm number and in transition zone length (Figure 4A and 5B). WAPL-1 protein localization functions as a marker of progenitor zone (mitotic region) (Crawley *et al*. 2016; Kocsisova *et al*. 2018). The *mir-235; mir-58.1 mir-83 him-8; mir-4807-4810.1* mutant male germlines displayed slower meiotic progression compared to control (Figure 6A). With a chase time of 10 hours, 57% of control male germlines had EdU-labelled spermatids, compared to 19% in the *mir-235; mir-58.1 mir-83 him-8; mir-4807-4810.1* mutants (Figure 6B). EdU-labeled cells were observed at a more proximal location in the control males (Figure 6C), indicating that the rate of meiotic progression in the mutants is slower than control males. There was no difference in the size of the EdU labelled region between control and mutant males immediately following the EdU pulse at 0 hours (Figure S6E), suggesting comparable rates of mitosis in the progenitor zone. Together, we found a slower rate of meiotic progression in the *mir-235; mir-58.1 mir-83 him-8; mir-4807-4810.1* mutant male germlines, which could lead to the observed slower rate of sperm production.

**Figure 6.**
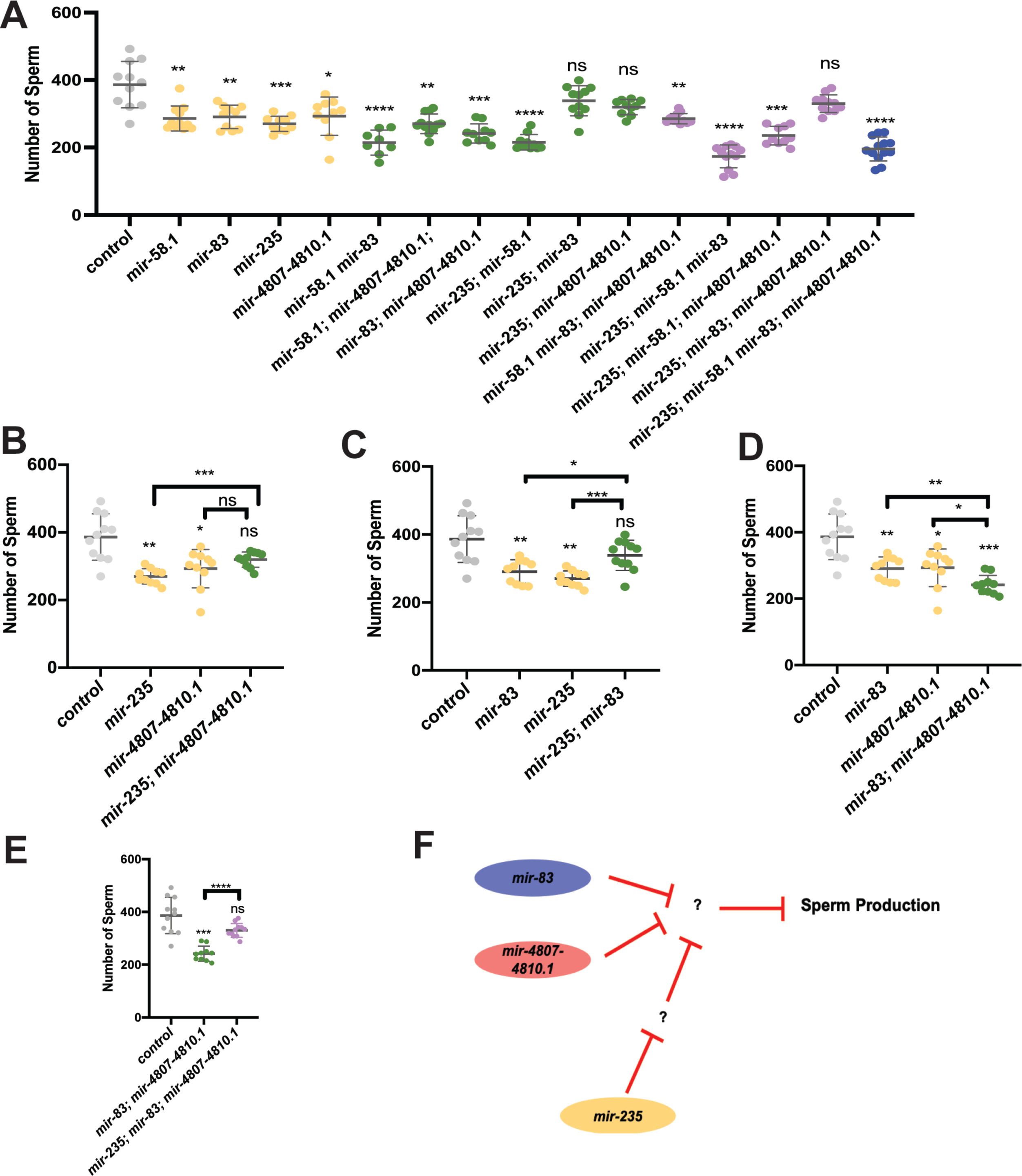
miRNA mutant males displayed slower meiotic progression. Control and *mir-235; mir-58.1 mir-83 him-8;mir-4807-4810.1* males were fed with EdU-labeled bacteria for 2.5 hours (pulse) to label cells in S phase, then transferred to unlabeled bacteria for 10 hours (chase) to assess the movement of these EdU+ cells. (A) Representative fluorescence images of male control or mutant gonads stained for EdU (green, left), WAPL-1 (red, middle), and DAPI (blue, right). White line indicates proximal boundary of progenitor zone (PZ), transition zone, pachytene, and condensation zone (diplotene+karysome+diakinesis). White arrows indicate the most proximal end of EdU+ cells. Red arrows indicate representative spermatids. Scale bars: 10µm. (B) The meiotic stage of the most proximal EdU+ cells was determined. The percentage of germlines in spermatogenesis, condensation zone or pachytene is shown. n=37. (C)The distance traveled by the most proximal EdU+ cells after chase of 10 hours. Each data point indicates the cell diameter (c.d.) of most proximal EdU+ cells from the edge of the PZ in individual gonads. Error bar indicates mean ± SD. Statistical test between control and mutant was performed by t-test with p<0.0001.

Next, we investigated whether the *mir-235; mir-58.1 mir-83 him-8; mir-4807-4810.1* mutant hermaphrodites also displayed a slower rate of meiotic progression during oogenesis. The brood size was reduced in the *mir-235; mir-58.1 mir-83 him-8; mir-4807-4810.1* mutants (Figure S7A) compared to controls. We analyzed meiotic progression in control and mutant hermaphrodites after a 4-hour EdU pulse followed by a 20 hour chase. First, the transition zone length was not affected (Figure S7B) unlike what was observed in the male germline (Figure 5B). Further, the distance of the most proximal EdU-labeled cells from the edge of progenitor zone is shorter in the mutants immediately following the EdU pulse at 0 hours and after the 20 hour chase (Figure S7C,D), which indicates a slower rate of EdU incorporation in the *mir-235; mir-58.1 mir-83 him-8;mir-4807-4810.1* mutant hermaphrodites, while meiotic progression was largely not affected because the percentage of germlines in either pachytene or condensation zone (diplotene, diakinesis, and oocyte) in *mir-235; mir-58.1 mir-83 him-8;mir-4807-4810.1* mutants was comparable to control hermaphrodites (Figure S7E). The slower rate of EdU incorporation could be due to a slower rate of mitosis or meiotic entry.

### Computational prediction of miRNA-target network for male gonad-enriched miRNAs involved in sperm production

To begin to understand the network of target mRNAs for *mir-58.1*, *mir-83, mir-235,* and *mir-4807-4810.1* in the process of sperm production, we performed computational analysis of the set of predicted targets using the Targetscan algorithm (Jan *et al*. 2011). Although the decreased sperm production in these miRNA loss of function mutants may be caused by disruption of miRNA function in somatic cells, we focused our computational analysis on the potential regulatory roles for miRNAs in the germline. We further filtered the list of predicted targets to focus on miRNA target mRNAs that are present in the *C. elegans* germline using published transcriptome data (Ortiz *et al*. 2014; Tzur *et al*. 2018). Gene ontology analysis with DAVID (Sherman *et al*. 2022) revealed an enrichment of target mRNAs categorized as genes associated with biological processes such as cell division, meiotic cell cycle, chromatin organization, mRNA processing (Table S6). KEGG pathway analysis of predicted germline mRNA targets indicated the possible regulation of pathways involving Notch signaling, RNA degradation, and MAPK by *mir-235*, *mir-4807-4810.1*, *mir-58.1*, and *mir-83* (Table S7). Additionally, network visualization revealed that many targets are potentially shared by *mir-235*, *mir-4807-4810.1*, *mir-58.1*, and *mir-83* (Figure S8). Together, this suggests genetic interactions between these miRNAs could likely be mediated through shared targets or pathways.

## Discussion

Here we identify a set of male gonad-enriched miRNAs that function to regulate fertility and fecundity in males and hermaphrodites. Three miRNAs were necessary for optimal male mating, *mir-83, mir-789.2,* and *mir-2221. mir-83* mutant males showed few interactions with hermaphrodites, suggesting defects in the ability to detect or respond to hermaphrodites. Three miRNAs, *mir-58.1, mir-83,* and *mir-235,* and one miRNA cluster, *mir-4807-4810.1* were found to be necessary for the regulation of sperm production under normal growth conditions, possibly through the regulation of meiotic progression in the early stages of prophase I. An additional three miRNAs, *mir-49, mir-57,* and *mir-261,* and one miRNA cluster, *mir-357/358,* were found to be necessary for sperm production under elevated temperature conditions. While the mechanism for this temperature sensitivity is unknown, mutant phenotypes for other miRNA loss of function alleles have also only been observed in conditions of a sensitized background or environmental stress (Ambros and Ruvkun 2018). Genetic analysis suggests a complex regulatory network with both parallel and antagonistic activity for male gonad-enriched miRNAs. Together, these data indicate male gonad-enriched miRNAs are necessary in males and hermaphrodites for optimal production of sperm.

### New functions identified for male gonad-enriched miRNAs

The *C. elegans* germline shows a dynamic regulation of gene expression to allow for the proliferation and differentiation of germ cells and the production of functional gametes (Reinke *et al*. 2000, 2004; Ortiz *et al*. 2014; Tzur *et al*. 2018; Ebbing *et al*. 2018). Translational regulation through the 3’ UTR is pervasive in the germline (Merritt *et al*. 2008) and miRNAs have been shown to be expressed and, with their associated Argonaute proteins, to contribute to the regulation of germ cell development in *C. elegans* (Bukhari *et al*. 2012; Lehrbach *et al*. 2012; Dallaire and Simard 2016; McEwen *et al*. 2016; Brown *et al*. 2017; Diag *et al*. 2018; Minogue *et al*. 2018; Bezler *et al*. 2019). Small RNA sequencing of isolated gonad arms from males and hermaphrodites was first described by Bezler *et al* (2019) and results presented here are consistent with this study. Based on our differential miRNA expression pattern between male and hermaphrodite gonads, a subset of 29 miRNAs were identified that were enriched in male gonads relative to hermaphrodite gonads.

We have defined new functions of *mir-58.1*, *mir-83, mir-235*, and *mir-4807-4810.1* in the regulation of sperm production in normal growth conditions. Additional miRNAs were found to be necessary only in conditions of elevated temperature. Because the miRNA mutant males analyzed herein are fertile, these functions of miRNAs likely contribute to the robustness and fidelity of sperm production rather than function as essential regulators of the core machinery of spermatogenesis.

Previous studies have characterized functions for *mir-58, mir-83,* and *mir-235* in *C. elegans.* The *mir-58* family functions to regulate the TGF-ß pathway to influence growth and dauer formation (de Lucas *et al*. 2015) and to prevent apoptosis during embryogenesis (Sherrard *et al*. 2017). *mir-83* modulates the migration of distal tip cells particularly in response to temperature stress (Burke *et al*. 2015), and coordinates autophagy with aging (Zhou *et al*. 2019). Lastly, *mir-235* has been shown to keep neural progenitor cells in a quiescent state (Kasuga *et al*. 2013; Kume *et al*. 2019), to mediate dietary restriction-induced longevity (Xu *et al*. 2019), and to protect the worm from graphene oxide toxicity in intestine (Guo *et al*. 2020). These miRNA functions may be independent of, or have an indirect connection to, the phenotypes we observe in sperm production. For example, loss of *mir-58.1* could result in mis-regulation of the TGF-ß pathway in the soma or the germline leading to changes in gene expression that could indirectly affect sperm production. Further analysis to identify direct targets and pathways is required to determine the mechanism of action of these male gonad-enriched miRNAs and it is not yet known if these miRNAs function in the germline, somatic gonad, or in other somatic cells.

The alleles for the *mir-4807-4810.1* cluster delete a 3.4kb region of the *Y59E1B.1* gene, so it is possible that it is the loss of Y59E1B.1 that causes the lower number of sperm in *mir-4807-4810.1* mutants. However, loss of both clusters of *mir-4807-4810.1* (3.4kb deletion) and *mir-1018-4923*.1 (457bp deletion), didn’t result in a reduction in sperm number, despite the loss of additional *Y59E1B.1* sequence. Further, *Y59E1B.1* is not expressed in hermaphrodite or male gonads (Bezler *et al*. 2019; Tzur *et al*. 2018). Together, the reduced sperm count in *mir-4807-4810.1* mutant males is not likely to be caused by the disruption of *Y59E1B.1* expression.

### Four male gonad-enriched miRNAs may regulate meiotic progression

Of the four miRNAs found to regulate spermatogenesis, two were observed to be necessary in both males and hermaphrodites, *mir-58.1,* and *mir-235,* and two were observed to be necessary only in males, *mir-83* and the miRNA cluster *mir-4807-4810.1.* While the onset of sperm production is modestly delayed in *mir-4807-4810.1,* it is unaffected in the other three mutants. However, all four miRNAs are observed to be required for the normal rate of sperm production. Loss of *mir-58.1*, *mir-83, mir-235*, and *mir-4807-4810.1* in males did not affect the rate of mitosis but did result in a slower rate of meiotic progression, which could account for the reduced number of sperm observed after the L4 molt stage.

In addition, *mir-58.1*, *mir-83, mir-235*, and *mir-4807-4810.1* were found to be necessary for normal progression through the transition zone. Surprisingly, despite the slower rate of meiotic progression, the transition zone was observed to be shorter rather than longer in single or multiply mutant worms. The transition zone in the *C. elegans* gonad is the region of the gonad in which germ cells first enter meiosis and show the polarized, crescent shaped nuclear morphology that corresponds with the early chromosome pairing events in leptotene/zygotene of prophase I (Dernburg *et al*. 1998; Colaiácovo 2013). In wild-type animals, germ cells lose this crescent morphology as they complete chromosome pairing and progress to pachytene. Mutant hermaphrodites that have defects in chromosome pairing and synapsis, such as *syp-1*, display an extended transition zone region with more polarized nuclei (MacQueen *et al*. 2002). In contrast, *plk-2* mutant hermaphrodites have defects in the synapsis checkpoint regulation and display a shorter transition zone region despite showing asynapsis of chromosomes (Harper *et al*. 2011). We hypothesize that the shorter transition zones observed in *mir-58.1, mir-83, mir-235,* and *mir-4807-4810.1* mutants may indicate that this set of miRNAs promotes the synapsis checkpoint (Jaramillo-Lambert *et al*. 2007; Harper *et al*. 2011). This could result in germ cells exiting leptotene/zygotene with asynapsis of chromosomes possibly causing delays or defects in meiotic progression to form haploid spermatids in males. Interestingly, the Targetscan algorithm identifies one binding site in the *syp-1* 3’UTR for the *mir-49/mir-83* miRNA family. One model is that the shorter transition zone in *mir-83* mutants could be due in part to mis-regulation of *syp-1.* Future work could examine *syp-1* and other predicted direct targets of *mir-58.1, mir-83, mir-235,* and *mir-4807-4810.1* with meiotic regulatory roles, such as *htz-1*, and *dpy-28* (Figure S8), suggested by germline transcriptome gene ontology analysis (Ortiz *et al*. 2014). Alternatively, a shorter transition zone may reflect accelerated age-dependent changes in the germline (Kocsisova *et al*. 2019), though no other gross morphological changes indicating accelerated aging were observed.

### miRNAs function in complex genetic networks in male gonads

A simple model of the male gonad-enriched miRNAs that are involved in male mating or sperm production is that these miRNAs function together to regulate shared pathways. Such functional redundancy has been observed for miRNA family members, which share a common 5’ “seed” sequence (nucleotides 2-7) (Abbott *et al*. 2005; Alvarez-Saavedra and Horvitz 2010; Duchaine and Fabian 2019). In our set of male gonad-enriched miRNAs, *mir-49* and *mir-83* are in the same miRNA family, as are *mir-789.2* and *mir-789.1*. In this work, additive effects of missing multiple family members for these two families were not observed in mating assays or sperm quantification analysis. Surprisingly, *mir-83; mir-49* and *mir-789.1; mir-789.2* double mutant males displayed partially suppressed mating defects compared to *mir-83* and *mir-789.1* single mutant males, while the sperm production defect observed in *mir-83* was suppressed by loss of *mir-49* at 20°C, but not at 25°C. These data indicate distinct roles of miRNA family members. This may reflect target recognition driven by the differences in the 3’ sequences between miRNA family members (Chipman and Pasquinelli 2019). In addition, these miRNAs may show differences in their spatial or temporal expression patterns driving differences in target and pathway regulation. Such overlapping and non-overlapping targets for miRNA family members is observed in the *let-7* family in *C. elegans* (Abbott *et al*. 2005; Broughton *et al*. 2016).

The set of gonad-enriched miRNAs involved in sperm production, *mir-58.1, mir-83, mir-235,* and *mir-4807-4810.1,* are not all in the same miRNA family and thus are not necessarily predicted to regulate common targets. Overall, analysis of multiply mutant worms indicated a trend that *mir-58.1, mir-83, mir-235,* and *mir-4807-4810.1* double, triple, and the quadruple mutant had stronger defects than the set of single mutants. However, there were exceptions, which suggest more complex genetic relationships. Our data are not consistent with a simple additive model for miRNA function but rather indicate that these male gonad-enriched miRNAs have targets and pathways that can act antagonistically. For example, *mir-235* shows opposing activity to *mir-83* in double mutant strains but shows additive activity with *mir-58.1.* Because miRNAs typically function as negative regulators of their downstream targets, this antagonism is expected to reflect the indirect effects of target mis-regulation rather than opposing activities on direct shared targets.

### Male gonad-enriched miRNAs may buffer environmental stress

In addition to *mir-58.1, mir-83, mir-235,* and *mir-4807-4810.1*, five miRNAs were identified to promote sperm production in conditions of moderate temperature stress (25°C). This is consistent with the model that miRNAs function to buffer environmental stressors possibly by acting as fine tuners of gene expression (Burke *et al*. 2015; Isik *et al*. 2016; Tran *et al*. 2019; Guo *et al*. 2020; Pagliuso *et al*. 2021). Thus, miRNA function may only be revealed under stressful or sensitized conditions (Brenner *et al*. 2010). For example, some male gonad-enriched miRNAs are mis-regulated in worms exposed to graphene oxide, which is toxic to the germline (Zhao *et al*. 2016), including *mir-2210*, *mir-4810*, and *mir-4807*. Our results highlight a role of miRNA in regulating sperm production in face of moderate temperature stress and this conditional role of miRNAs may be observed under other stress conditions. It is important to note that this elevated temperature of 25°C is within the normal cultivation temperature range of *C. elegans* through which wild-type male and hermaphrodite worms maintain fertility.

Taken together, we have identified a set of male gonad-enriched miRNAs that are necessary for normal sperm production, in part through the regulation of meiotic progression. Genetic data indicates a complex network of miRNAs in the male gonad, which will require a comprehensive analysis of the set of miRNA targets to elucidate.

## Materials and Methods

### Strains

All *C. elegans* strains were maintained by growing on AMA1004 (Casadaban *et al*. 1983) seeded NGM plates. As a control, sperm quantification in control and *mir-235; mir-58.1 mir-83 him-8; mir-4807-4810.1* mutant males was also performed following growth on OP50 seeded plates (Figure S4D). To facilitate the phenotypic analysis on both males and hermaphrodites, *him-8(e1489)* and *Phis-72::HIS-72::GFP(stls10027)* were crossed in to all miRNA mutant strains to facilitate the sperm quantification in males (Huang *et al*. 2012). This is referred to as “*him-8; his-72::gfp”* herein. A strain with *him-8; his-72::gfp* was used as a control in experiments unless otherwise specified. All strains analyzed in this paper are listed in Table S8. The UY264 strain with *mir-49(zen99)* was a gift from Dr. Anna Zinovyeva. All *n* alleles of miRNA genes were described in Miska *et al* (Miska *et al*. 2007) and were outcrossed 4x. The *mir-57(gk175)* allele was generated by the *C. elegans* Deletion Mutant Consortium (The *C. elegans* Deletion Mutant Consortium 2012).

### CRISPR/Cas9 mutagenesis

miRNA mutants were generated using CRISPR/Cas9, by which miRNA sequences were knocked out when Cas9-mediated cuts with a single sgRNA or two sgRNA sites were repaired with a template missing the mature miRNA sequence (Dickinson *et al*. 2013). Briefly, sgRNA encoding sequence and self-excising cassette (SEC)-containing repair templates were assembled in a plasmid using SapTrap (Dickinson *et al*. 2015, 2018). The sgRNA, SEC, and repair template plasmid, Cas9 expression plasmid, and co-injection markers (pGH8, pCFJ104, and pCFJ90) were injected into the gonads of young adult hermaphrodites. When two sgRNAs were used, the second sgRNA was cloned to Cas9-containing plasmid pDD162 with Q5 mutagenesis kit. Plasmids were obtained from Addgene. Candidates were selected using the dominant Roller phenotype and hygromycin resistance. The SEC was removed by heat shock and the candidates were sequenced to confirm accurate genome modification. All miRNA loss of function mutants constructed by CRISPR/Cas9 were backcrossed to N2 twice before phenotypic analysis. Strains with new mutant alleles and *him-8; his-72::gfp* were then constructed. The new miRNA loss of function alleles generated in this paper and the designs for CRISPR are listed in Table S9.

### Small RNA sequencing

At Day 0, L4 stage males or hermaphrodites were picked from N2 plate. At Day 1, young adult worms were picked to 1xPBS with 1mM levamisole to immobilize them. Two fine-gauge needles were used to release the gonads and separate them from the body. A single gonad arm was isolated from individual hermaphrodites, and the spermatheca was removed. For each sample, 300 gonads from males or hermaphrodites were collected in Trizol (Invitrogen 15596026) and stored in the -80°C freezer prior to total RNA prep (DirectZol RNA microprep kit, Zymo R2060). Three independent biological replicate samples were prepared for N2 males and N2 hermaphrodites. The RNA samples were processed in University of Wisconsin-Madison Biotechnology Center Gene expression Center and DNA Sequencing Facility for TruSeq Small RNA Library construction and sequencing. The RNA-seq analysis was performed on Galaxy platform. Adapters of small RNA seq reads were trimmed with Triommatic, including (1) RNA 3’ Adapter: TGGAATTCTCGGGTGCCAAGG; (2) PCR_Primer Index: CAAGCAGAAGACGGCATACGAGAT; (3) RNA_PCR_Primer: AATGATACGGCGACCACCGAGA; (4) PCR_Primer_Index: CAAGCAGAAGACGGCATACG. Reads were aligned to the reference genome assembly Ce10 with Bowtie2. The aligned reads were annotated with miRbase22 with htseq-count. Differential expression analysis was performed with DESeq2. miRNAs were considered significantly expressed between male and hermaphrodite if they had log2(fold change) > 1 and P-adj < 0.01.

### Assays for male fertility and fecundity

#### Sperm quantification

A single male was picked to a 3µL drop of sperm buffer (50mM HEPES pH7, 25mM KCL, 45mM NaCl, 1mM MgSO4, 5mM CaCl2, 10mM Dextrose pH7.8) on a glass coverslip, which was placed directly on a slide allowing the release of sperm from the worm. The number of HIS-72::GFP positive sperm was counted using epifluorescence microscope at 40X.

#### Sperm onset

Early L4 stage males of control and mutant strains were observed for presence of haploid spermatids using the *his-72::gfp* transgene expression to detect individual sperm with characteristic condensed chromatin. Early L4 stage males were further staged based on whether tail retraction was observed (Nguyen *et al*. 1999). The percentage of males with spermatids was calculated for all strains.

### Assays for hermaphrodite fertility and fecundity

#### Brood size Assays

At Day 1, L4 stage hermaphrodites were picked from control or mutant strains, and were transferred to a new plate every day until production of progeny ceased. The number of progeny was counted on each plate and the total number of progeny was calculated. Unhatched eggs were not scored in this assay. **Brood size after mating** At Day 0, matings were set up with one mutant or control hermaphrodite and 1-5 control males to ensure successful mating and transfer of sperm. On Day 1, males were removed from the plate, and the hermaphrodite was picked to a new plate. Successful mating was confirmed by the presence of ∼50% male cross progeny. The hermaphrodite was transferred to a new plate each day until production of progeny ceased. The number of progeny was counted on each plate and the total number of progeny was calculated.

#### DAPI-staining of isolated gonads

Gonads were dissected for DAPI staining (Gervaise and Arur 2016; Kocsisova *et al*. 2018). At Day 0, males at L4 stage were picked from control or mutant strains. At Day 1, young adult males were washed 3 times with M9.

On a watch glass, males were transferred to M9 with 1∼3µL 100mM levamisole. Two fine-gauge needles were used to cut the worms at the pharynx and release the gonad arm. Dissected worms were transferred to a glass conical tube with methanol stored at - 20°C for at least 1 hour or overnight. After methanol fixation, the worms were washed 3 times with 1xPBS-T, then transferred to 1mL glass tube. The worms were incubated in 200uL 1µg/mL DAPI solution in PBS-T in the dark for 30mins. After 3 washes with 1x PBS-T, the worms were transferred to an agarose pad with DABCO Mounting Medium (1,4-Diazobicyclo-(2,2,2) octane, glycerol, 1xPBS, pH 8.6). Alternatively, the Vectashield mounting medium with DAPI was used directly (Vector H1200). An eyelash pick was used to position the gonads, and extra fluid was removed with glass pipette. A glass coverslip (24X50) was placed gently on top of agarose pad and nail polish was applied to the edge of coverslip. The slides were stored at 4°C in the dark overnight. Images were captured with a Nikon Eclipse Ti Confocal microscope and images were analyzed using Nikon Elements software.

#### EdU pulse chase

In this study, EdU labeling was performed by feeding worms with EdU-labelled *E.coli*. The EdU pulse chase experiment was performed as previously described (Kocsisova *et al*. 2018). To prepare EdU-labeled bacteria, 4mL freshly overnight LB culture of *E.coli* strain MG1693, which carries a mutation in thyA, was added to liquid culture (5mL of 20% glucose, 10 mg/mL of thiamine, 120µl of 5mM thymine, 100µL of 1M MgSO4, 100µL of 10mM EdU, 100 mL of M9 buffer) for growth overnight. From the liquid culture, EdU-labeled *E.coli* was concentrated, and then plated on M9 agar petri dishes (M9 buffer + agar) to make EdU-labeled growth plates for worms. Gravid hermaphrodites from control or mutant worms were bleached to collect a synchronized population of L1s. L4 stage male and hermaphrodites were picked at around 41 hours, and 46 hours after seeding L1s on NGM, respectively. Then the worms were washed using M9 and then transferred to the bacterial lawn on the EdU-labeled plates. The EdU plates with males and hermaphrodites were incubated at 20°C for 2.5 hours, and 4 hours, respectively, after which, the worms were washed off h with M9. After the EdU pulse, worms were either dissected to release the gonad arm (chase0) or transferred to NGM plates. Males and hermaphrodites were grown at 20°C for duration of 10 hours and 20 hours, respectively, during which EdU-labeled cells progress proximally, and the worms were dissected to release the gonad arm after the 10- or 20-hour chase. The fixation and antibody staining were performed in glass tubes as previously described (Gervaise and Arur 2016; Kocsisova *et al*. 2018). Briefly, the isolated gonads were fixed with 3% paraformaldehyde (PFA) for 10mins first, then in cold methanol at -20°C for at least 1 hour to overnight. After washing with 1xPBS-T, gonads were blocked in 30%Normal Goat Serum (NGS) at room temperature for 1 hour. The gonads were then incubated with a primary antibody rabbit-anti-WAPL-1 (1:100 diluted in 30%NGS) for 4 hours to overnight at room temperature followed by incubation in a goat-anti-rabbit IgG-conjugated Alexa Fluor 594 secondary antibody solution (1:400, 2 hours at room temperature or overnight at 4 °C). Next, an EdU Click-iT reaction was performed to detect EdU according to manufacturer’s instructions (Click-iT EdU Alexa Fluor 488 Imaging Kit, Thermo Fisher Scientific). Gonads were incubated in the EdU cocktail mixture with 8.5µl of 10x buffer, 76.5µL of ultrapure water, 4µL of 100mM CuSO4, 0.25 µl of the 488 nm dye Azide, and 10µL of 1x buffer additive added in order for 30 mins at room temperature in dark. After washing with 1xPBS-T, the gonads were mounted on agarose pads with the Vectashield mounting medium with DAPI (Vector H1200). The slides were stored at 4°C in the dark overnight. Images were captured within 72 hours with a Nikon Eclipse Ti Confocal microscope at 60x. Images were analyzed using Nikon Elements software. The edge of progenitor zone (PZ) in the germline was determined by WAPL-1 and DAPI labeling, because WAPL-1 signal decreases as cells enter meiotic prophase (Crawley *et al*. 2016), and chromosome organization is different in PZ, transition zone, pachytene, and condensation zone (diplotene and diakinesis) with DAPI staining (Shakes *et al*. 2009).

## Data Analysis

All statistical analysis was performed using GraphPad Prism software. Details for each statistical test are found in the figure legends.

## Data Availability

Strains with miRNA deletion alleles and plasmids used for CRISPR-Cas9 genome modification are available upon request. The small RNA sequencing data discussed in this publication have been deposited in NCBI’s Gene Expression Omnibus and are accessible through GEO Series accession number GSE239800.

## Acknowledgments

The authors thank the University of Wisconsin-Madison Biotechnology Center Gene Expression Center & DNA Sequencing Facility for providing library preparation and next generation sequencing services and Phillip Ross and Trestle Biosciences for sequencing analysis and data visualization support. Some strains used in this study were obtained from the *Caenorhabditis* Genetics Center (CGC), which is funded by NIH Office of Research Infrastructure Programs (P40 OD010440). We thank Dr. Carmela Rios for assistance with CRISPR-Cas9 genome modifications to generate new mutant alleles, Dr. Tim Schedl for sharing the anti-WAPL-1 antibody and the MG1693 *E. coli* strain for EdU pulse chase experiments, and Dr. Anna Zinovyeva for sharing the UY264 *mir-49*(*zen99*) X strain.

## Supporting Information

## Supplemental Methods

### Supplemental Figure legends

**Figure S1. miRNA profile comparison between replicates in male and hermaphrodite.** Each miRNA identified was plotted in Log2 (normalized counts+1) to show the correlation between biological replicates in hermaphrodite (A) and male gonads (B). And Pearson correlation coefficients with 95% confidence intervals are shown on the top left corner. (C) Heat map showing the distance between the samples Male replicates and hermaphrodite replicates clustered into a group.

**Figure S2. miRNAs regulate male mating.** (A) Percentage of mating success was determined by the number of mating plates with *unc-17(e245)* hermaphrodites with presence of cross progeny. The number of successful matings and total number of mating assays is indicated above each bar. (B-C) Interaction between hermaphrodites and mutant males was further analyzed by sperm transfer assay. (B) Control and mutant males were analyzed for mating interactions with hermaphrodites. Bar graphs show % of control (*him-8; his-72*) or mutant males that interacted with hermaphrodites. (C) The percentage of the interactions in (B) that led to successful sperm transfer from control or mutant males was determined. The number of successful sperm transfers and total number of interactions is indicated above each bar. (D) miRNA family members for *mir-83* and *mir-789.1.* Seed sequence (red) is shared between *mir-49* and *mir-83*, and between *mir-789.1* and *mir-789.2*. (E) The percent mating success of single and double mutant males in mating assay. The statistical analysis between control and mutants was conducted using Fisher’s exact test and shown as p>0.05; *, p<0.05; **, p<0.01; ***, p<0.001; ****, p<0.0001.

**Figure S3. The number of sperm in miRNA mutants at 20°C or 25°C.** Sperm quantification was performed for individual control and miRNA multiply mutant strains using *his-72::gfp* to detect haploid spermatids. Each dot in the scatter plot represents the number of sperm in individual worm and lines represent mean ± SD for each strain. The male sperm was counted at L4molt+5hours unless specified. All strains assayed with *him-8(e1489); stls10027 (Phis-72::HIS-72::GFP)* in the genetic background. The statistical analysis results are represented as ns, p>0.05; *, p<0.05; **, p<0.01; ***, p<0.001; and ****, p<0.0001 with Dunnett’s T3. (A) The number of male sperm in control and mutant carrying independent *mir-4807-4810.1* loss of function allele (*xwDf15*). (B) The number of male sperm at 20°C. (C)Analysis of sperm number in miRNA cluster mutants. (D) The number of male sperm at 20°C for *mir-58.1* and *mir-83* family members. (E) The number of male sperm at 25°C for *mir-58.1* and *mir-83* family members.

**Figure S4. Genetic interactions between *mir-58.1*, *mir-235*, and *mir-83* to regulate male sperm production. (**A)(B)(C) Selected data sets from multiply mutant sperm analysis (Figure 4A) were shown to highlight interaction. Each dot in the scatter plot represents the sperm count in an individual worm and lines represent mean ± SD for each strain. The statistical analysis results are represented as ns, p>0.05; *, p<0.05; **, p<0.01; ***, p<0.001; and ****, p<0.0001. The comparison between control (*him-8; his-72*) and the miRNA mutant is shown above each data set as Figure 4A, and other comparison pair is indicated by the line above them by Welch’s T-test. (D)The sperm count of control and miRNA quadruple mutant males on different food source. The statistical analysis was performed with one-way ANOVA followed by Tukey’s multiple comparisons test.

**Figure S5. Brood size of miRNA multiply mutants.** Genetic interactions were analyzed between *mir-58.1, mir-83, mir-235* and *mir-4807-4810.1.* Brood size analysis was performed for individual control and miRNA multiply mutant strains. Each dot in the scatter plot represents the brood size in individual worms and lines represent mean ± SD for each strain. All strains assayed with *him-8; his-72* in the genetic background. The statistical analysis results are represented as ns, p>0.05; *, p<0.05; **, p<0.01; ***, p<0.001; and ****, p<0.0001. The comparison between control (*him-8; his-72*) and mutant is indicated above the data sets by Dunnett’s T3. (A) Brood sizes in controls (gray), single (yellow), double (green), triple (purple), and quadruple (blue) mutant hermaphrodites. (B-D) Selected datasets from data shown in (A) to highlight interactions in single and double mutants. The comparison between single and double mutants is indicated by the line above them with Welch’s T-test.

**Figure S6. Analysis of DAPI stained gonads in multiply mutant males and analysis of the timing of sperm onset in control and miRNA mutant males.** (A-B) Individual DAPI stained gonads were analyzed for the length of the mitotic zone (A) in miRNA single mutants (A) and multiply mutants (B), determined by the number of cell rows from the distal end to the start of the transition zone with nuclei with polarized chromatin in a crescent morphology. Each dot in the scatter bar represents the number of cell rows in mitotic region, error bar indicates mean ± SD; The statistical analysis between control (*him-8; his-72*) and mutant was conducted by Dunnett’s test. (C-D) Sperm onset was determined by the presence of haploid spermatids in early L4 before tail retraction (C) and in early L4s with tail retraction (D). The bar graphs represent the percentage of worms observed with spermatids, and the number of worms analyzed is indicated above each bar. The statistical test between control and mutant was conducted by Fisher’s exact test shown as ns, p>0.05; *, p<0.05; **, p<0.01. (E) The germlines of control and miRNA mutant males were labeled with EdU by feeding with EdU-labeled bacteria for 2.5 hours. Each dot represents the distance of the most proximal EdU+ cells from the edge of PZ (progenitor zone). The distance between control(*him-8*) and mutant male gonads were not significant by T-test.

**Figure S7. EdU pulse-chase analysis of multiply mutant hermaphrodites.** (A) Brood size was decreased in the miRNA quadruple mutant hermaphrodites. Each dot in the scatter plot represents the number of progenies in individual worms and error bar represents mean ± SD. (B)(C)(D) EdU labeling of S-phase cell in the germlines was achieved by feeding hermaphrodites with EdU-labelled bacteria for 4 hours(pulse), followed by transferred to unlabeled bacteria for 20 hours(chase). Hermaphrodite gonads were isolated after EdU labeling (chase0) or after chase (20 hours) for analysis when stained with DAPI, WAPL-1, and EdU. DAPI and WAPL-1 were used to define the proximal edge of PZ, transition zone, pachytene, and condensation zone (diplotene and diakinesis). Each dot represents one germline, and error bar indicates mean ± SD. The comparison between control and mutant was conducted by Welch’s t-test or unpaired t test. (B) Transition zone length of hermaphrodite germlines in cell diameter. n=10. (C) The distance of most proximal EdU+ cells from the edge of PZ after 4 hours of EdU labeling by feeding in cell diameter. n=12. (D) The EdU-labeled cells migrate proximally during 20 hours of chase. The distance was measured in cell diameter from the edge of PZ. (E) The meiotic stages of most proximal EdU+ cells. The bar graph represents percentage of germlines with the most proximal EdU+ cells in pachytene, and condensation zone. Fisher’s exact test was used to compare the meiotic stage profile between control and mutant after 20 hours of chase. n=12.

**Figure S8. *mir-235, mir-58.1, mir-235* and *mir-4807-4810.1* have potential shared targets.** miRNA-target network analysis of *mir-235, mir-58.1, mir-235* and *mir-4807-4810.1.* The miRNAs were shown in pentagon (red), and targets in ovals (light blue). For simplification, only shared targets were shown on the network.

**Table S1. Differential miRNA expression analysis between male and hermaphrodite gonad performed with DESeq2.**

**Table S2. Male fertility Analysis of miRNA mutants**

**Table S3. Brood size analysis of miRNA mutants**

**Table S4. Hermaphrodite sperm quantification of selected miRNA mutants**

**Table S5. Sperm quantification and brood size analysis of multiply miRNA mutants of *mir-58.1, mir-83, mir-235* and *mir-4807-4810.1***

**Table S6. Gene ontology analysis on predicted targets of *mir-58.1, mir-83, mir-235* and *mir-4807-4810.1***

**Table S7. KEGG pathway enrichment analysis on predicted targets of *mir-58.1, mir-83, mir-235* and *mir-4807-4810.1***

**Table S8. List of strains analyzed in this study**

**Table S9. New miRNA loss of function alleles generated in this study**

---

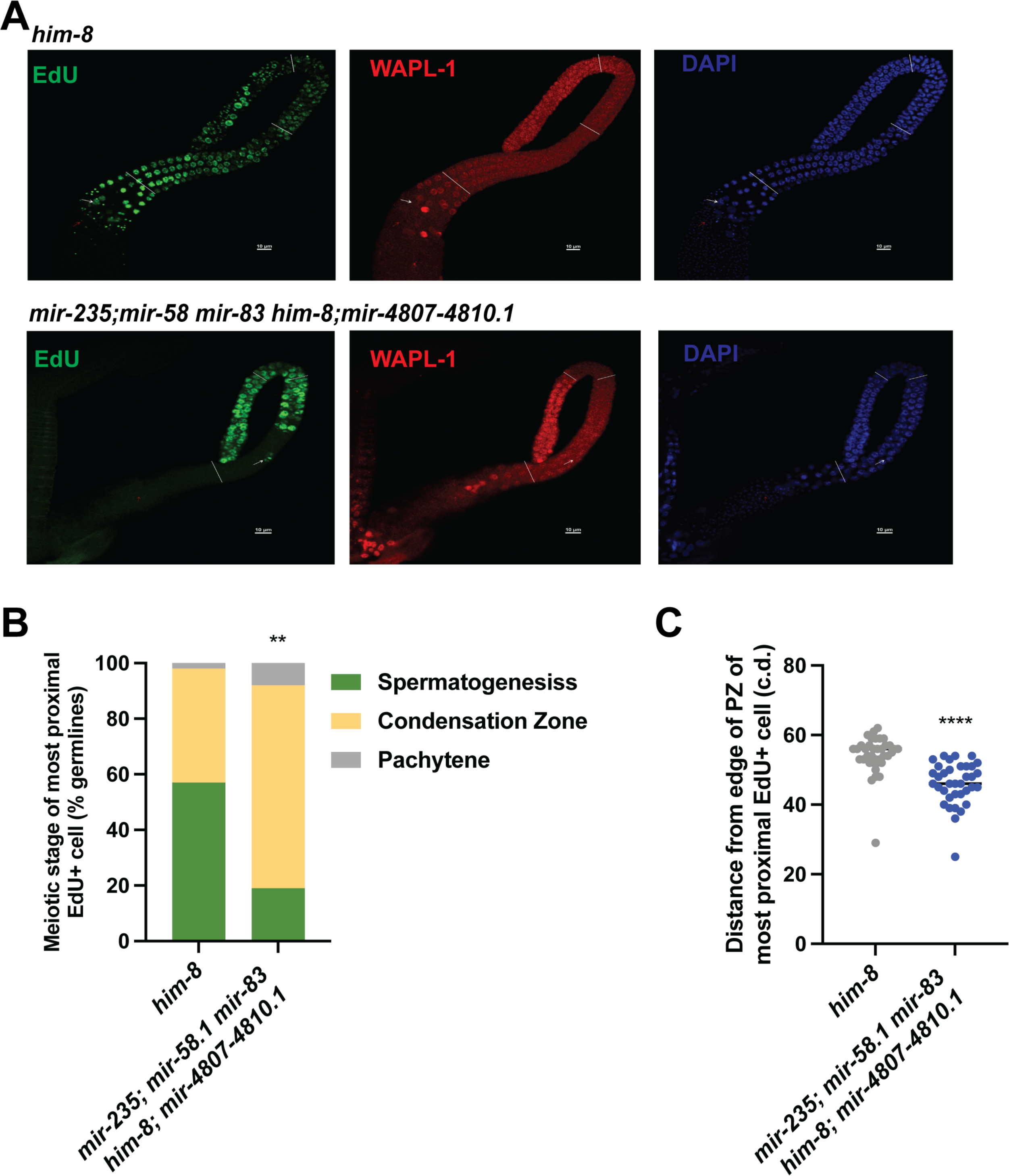

